# Mapping enzyme catalysis with metabolomic biosensing

**DOI:** 10.1101/2021.10.15.464597

**Authors:** Linfeng Xu, Kai-Chun Chang, Emory M. Payne, Cyrus Modavi, Leqian Liu, Claire M. Palmer, Nannan Tao, Hal S. Alper, Robert T. Kennedy, Dale S. Cornett, Adam R. Abate

## Abstract

Enzymes are represented across a vast space of protein sequences and structural forms and have activities that far exceed the best chemical catalysts; however, engineering them to have novel or enhanced activity is limited by technologies for sensing product formation. Here, we describe a general and scalable approach for characterizing enzyme activity that uses the metabolism of the host cell as a biosensor by which to infer product formation. Since different products consume different molecules in their synthesis, they perturb host metabolism in unique ways that can be measured by mass spectrometry. This provides a general way by which to sense product formation, to discover unexpected products and map the effects of mutagenesis.

## Introduction

Enzyme engineering uses an iterative cycle in which libraries of gene variants are designed, synthesized into proteins, and tested for the activity of interest^1^. The success of these engineering campaigns, however, is dependent on technologies for conducting these steps. Moreover, while there have been significant advances in design and build, test remains a bottleneck^2^. For example, the dominant strategy is well plate screening, because it is simple and flexible, and allows a variety of measurement techniques to directly quantify the product of the enzyme^2,3^. Well plate screening, however, is severely limited in scalability, testing just hundreds of variants per cycle^3^; this is a major issue because the likelihood of identifying superior variants scales with the number screened. Thus, alternative approaches based on selections, flow cytometry, and droplet microfluidics are valuable because they afford much higher throughput, screening >10^7^ variants per cycle^2,4^. However, a major constraint of these approaches is that they do not directly detect product formation, requiring a secondary assay to tether it to a detectable readout^4^. Identifying such assays can be challenging and, often, is not possible for enzymes of interest^2^. To enhance our ability to engineer enzymes through screening, a new method is needed that combines the generality of well plates with the scalability of microfluidics.

In this paper, we describe a screening approach that combines the scalability of microfluidics with the generalizability of mass spectrometry (MS) (**Figure 1**). Our approach leverages the background metabolism of the host as a biosensor with which to assess the activity of an enzyme embedded in it, and a novel microscale mass spectrometry technology to characterize metabolomic changes. Since the enzyme catalyzes molecules of central metabolism, its activity perturbs the host’s metabolite profile^1,5^, generating a signature that can be detected even if the enzyme product is not directly observed. Our approach thus provides a general way to map the catalysis of a mutated enzyme, to characterize the range of products it generates, and recover the sequences of variants with desired activities.

**Fig. 1.**
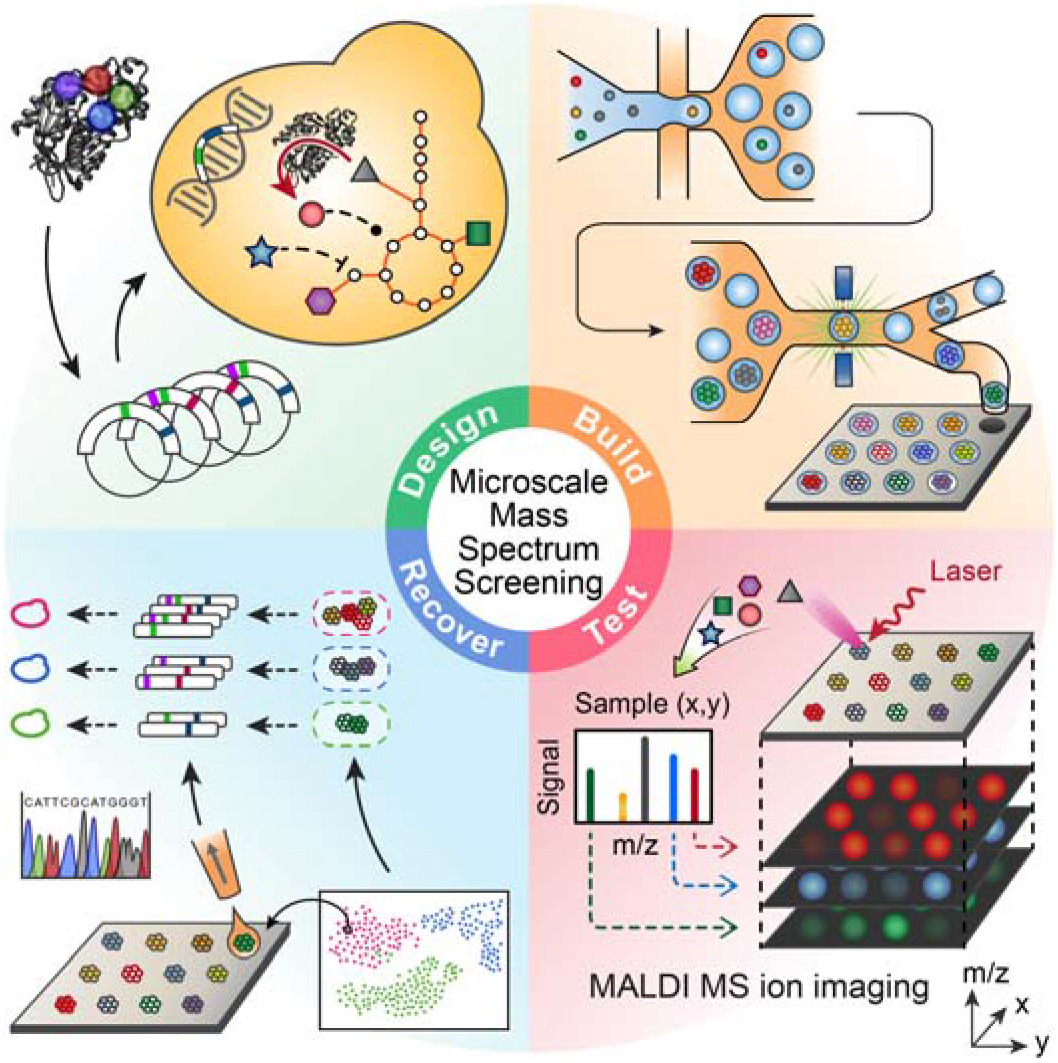
Metabolomic biosensing with microscale mass spectrometry provides a general strategy for screening enzymes. Enzyme variants are designed and transformed into yeast (design), then synthesized in the yeast where they consume molecules of central metabolism to generate product (build). Using printed droplet microfluidics, they are dispensed to a picoliter well array and subjected to MALDI-MS imaging to quantify cell metabolites (test). UMAP clusters cells according to metabolomic profile, where each cluster indicates a different enzyme phenotype. Desired mutants are extracted from the plate, sequenced, and confirmed in bulk cultures.

## Results

Matrix-assisted laser desorption ionization (MALDI) MS has been used for single cell metabolomics^6^ and to identify enzyme products from microbial colonies^7^. Here, we combine this approach with printed droplet microfluidics (PDM) to prepare, print, and screen all mutants from a semi-rationally varied 4-position library of the *Gerbera hybrida* G2PS1 type-3 polyketide synthase (t3PKS)^8^, comprising 1960 codon shuffled members (**Figure 2A**) (**methods**). This enzyme is responsible for biosynthesis of triacetic acid lactone (TAL) through condensation of a starter acetyl-CoA unit with two malonyl-CoA molecules and subsequent cyclization of the triketide chain ^8,9^. TAL has been used as a platform precursor for synthesis of high value chemicals commonly derived from fossil fuels. Mutations in the active site of these enzymes can alter the kinetics and spectrum of polyketide products formed^8,10^, potentially accessing novel products (**Figure 2B, Figure S1**). We synthesize and transform our library into *Yarrowia lipolytica*, encapsulating and culturing single cells in 300 pL droplets to generate isogenic colonies (**Figure S2**). Culture expansion produces additional material compared to a single cell, boosting MS signal and providing accurate metabolomic data.

**Fig. 2.**
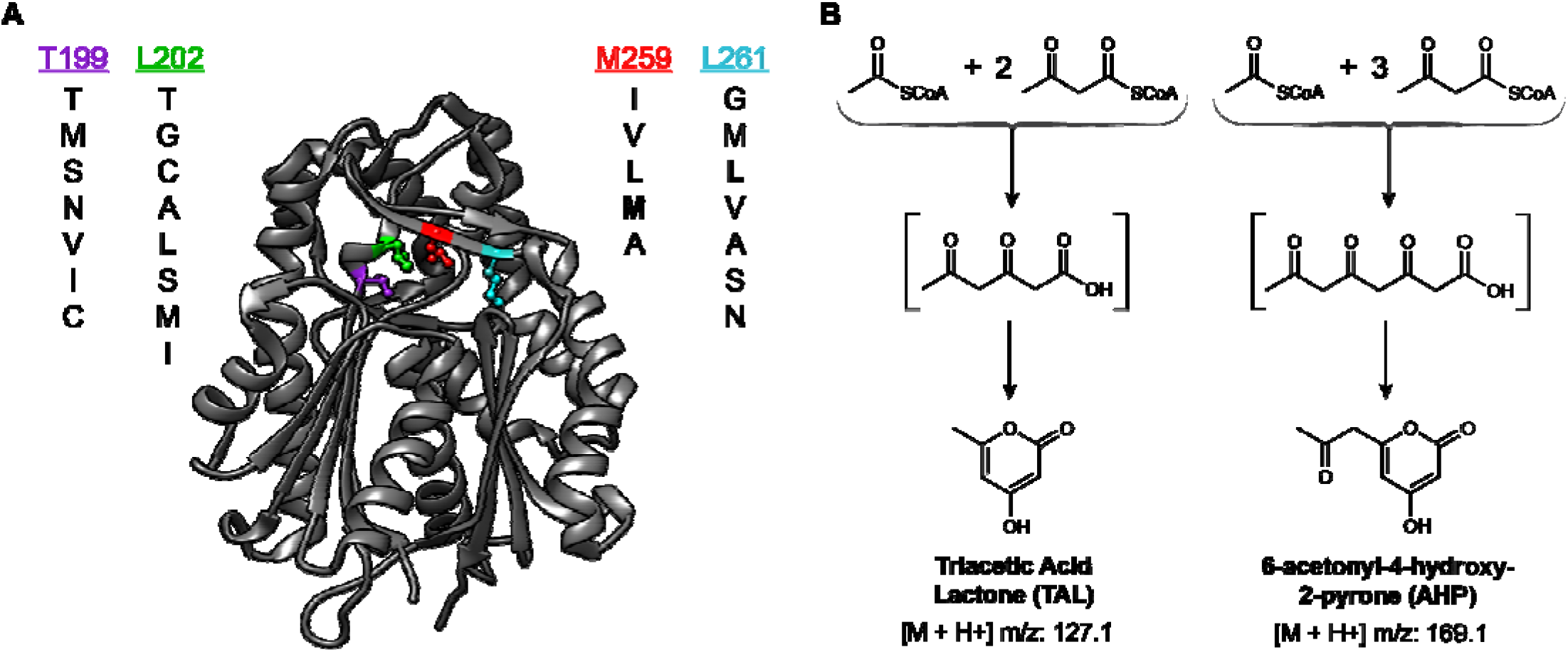
Overview of the type 3 PKS variant library. **A**, Crystal structure of *Gerbera hybrida* G2PS1 (PDB ID: 1EE0) showing the location of the four residues selected for mutagenesis and identity of the residue mutants. All residues except T199 directly form the active site cavity. **B**, The smallest condensation/cyclization products expected from type III polyketide synthase activity: Triacetic Acid Lactone (TAL, the native product of G2PS1) from one acetyl-CoA and two Malonyl-CoA and 6-Acetonyl-4-Hydroxy-2-Pyrone (AHP) from one acetyl-CoA and three Malonyl-CoA. Higher order polyketides, not shown, are possible from additional condensations of Malonyl-CoA.

To perform microscale MS (μMS), we use a high-density plate comprising a glass slide etched with 10,000 wells, having 80 μm diameters and rounded bottoms (**Figure. 3A**); this shape concentrates the material to the center (**Figure S3**), enabling accurate μMS quantitation. Higher capacity slides with the dimensions of a MALDI plate can accommodate 100,000 wells (**Figure S4**). To maximize throughput, all wells must be loaded with one colony, which we accomplish with printed droplet microfluidics (PDM)^11^ in ∼30 min (**Figure S5**). During printing, we scan the colonies, dispensing only ones falling within a narrow cell density range (**methods**)^11^.

**Fig. 3.**
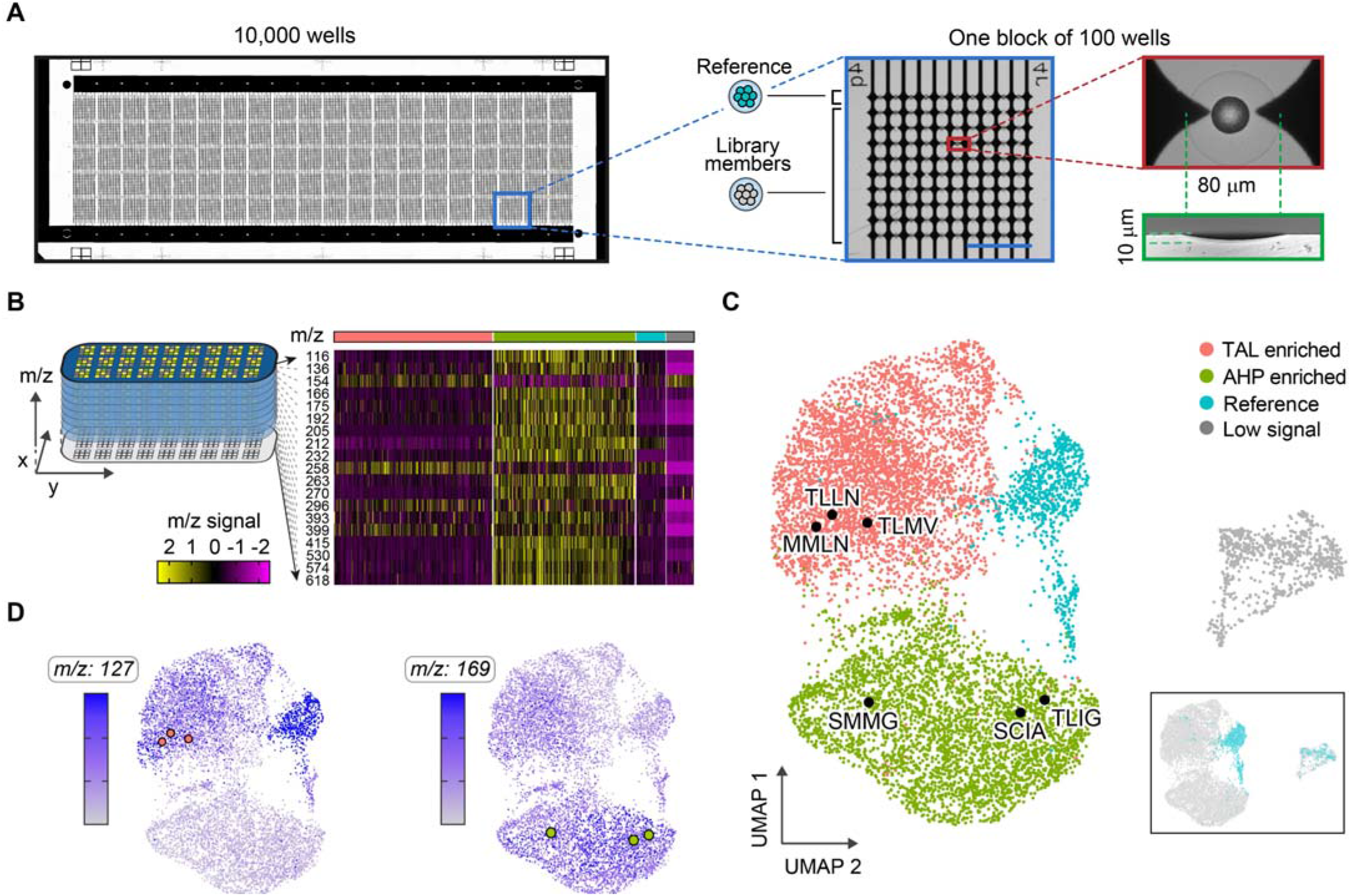
Mapping g2ps1 mutant catalysis through metabolic biosensing. **a**, Overview of slide (75 mm x 25 mm) for single variant μMS comprising 10,000 wells grouped in 100 blocks of 100 wells (magnified blue box, scale bar 1 mm. Each circular well is paired with electrodes for printed droplet trapping (red box) and has a rounded profile that concentrates desiccated metabolites to the center (green). **b**, μMS generates a spatial image of the substrate in which each pixel contains a full m/z spectrum. Well spectra are extracted and clustered into four groups using UMAP, **c**. Mutants of interest are recovered, sequenced, and overlaid on the clusters as black dots. Inset shows the reference strain based on known print locations. **d**, Heat maps for m/z 127 and 189 indicate heigh production of TAL in the upper and reference clusters, and high production of AHP in the lower cluster.

Using this system, we print 9,000 library members and 1,000 reference strains to designated positions. Once loaded, the plate is dried, spray-coated with matrix, and subjected to MALDI-MS imaging (**methods**), providing data on all detectable metabolites from m/z 30 to 630. The results are reported as an ‘image’ in which each pixel comprises a 600-dimensional vector of signal amplitude for each mass-to-charge ratio (m/z) (**Figure S6**-**S8, Table S1**). Thus, the data can be thought of as representing a 600 ‘color’ image, in which each slice reports the amplitude of the metabolite corresponding to the respective m/z. Importantly, MALDI’s soft ionization preserves DNA^12^, allowing PCR recovery of enzyme genes after MS analysis (**Figure S9**).

A unique and valuable property of μMS is that it provides information on many molecules in the host cells, enabling discovery of unexpected enzyme activities. A general way to identify activities is to plot alterations in the cell metabolome as a uniform manifold approximation and projection (UMAP), a dimensionality reduction technique used in single cell sequencing^13,14^ and MS imaging^15^ that projects high dimensional data onto a plane while preserving cluster information. To obtain the clearest UMAP clustering, we apply an algorithm to select the best m/z peaks for inclusion, comprising ∼60% of the total m/z intensity (**Figure 3B, Figure S9, S10, Movie S1, S2**) (**methods**). From this, we observe four clusters (**Figure 3C**) which, visually, resemble UMAP clusters of single cell RNAseq profiles, except they correspond to metabolite profiles^7,16^. The compact blue cluster is the 1000 control reference strains, which we confirm by substrate location (**Figure 3C**, *inset*). The gray island corresponds to wells with few cells that were mis-printed (**Figure S11**). This leaves two large clusters (red and green) which, presumably, correspond to cells with distinct metabolomic profiles. Since only the embedded enzyme varies between these cells, this implies the codon shuffled library exhibits two major activities that perturb cell metabolism in distinct ways. Mapping the m/z 127 data onto the clusters shows that the upper red island represents productive TAL mutants (**Figure 3D**, *left UMAP*). In addition to the desired product, some mutants generate another product dominantly produced in the lower green cluster. Upon careful inspection of the MS data, we determine that a molecule with the m/z value of 169 has differential abundance between these clusters (**Figure S12, Table S1, S2**). Using HPLC with tandem MS (HPLC-MS/MS), we confirm that the mass corresponds to the reported alternative product of this enzyme, tetraketide 6-acetonyl-4-hydroxy-2-pyrone (AHP)^10,17^. When we overlay the AHP amplitude (m/z 169) on the UMAP, it is predominantly found in the green cluster (**Figure 3D**, *right UMAP*).

To characterize how these disparate activities relate to enzyme sequence, we select and sequence ∼50 mutants from each cluster (**Table S3, S4**). Plotting the mutants versus normalized TAL and AHP shows colonies with extreme activity and, generally, that mutants efficient at producing one product are inefficient at producing the other (**Figure 4A**). To validate these results, we re-transform and test several mutants in bulk, finding good agreement (**Figures 4B, Figure S13, Table S3**). Interestingly, the TLLL mutant based on consensus design^18^ of TAL enriched members from the UMAP also shows high TAL production, though it retains higher AHP side-activity than the TLLN mutant (**Figure S14**). These results demonstrate that generation of these products can be inferred from clustering the host cells’ metabolomic profiles, even though the peaks associated with these molecules (m/z 127 and 169) are not included in the clustering. In this way, the host cell’s metabolite profile affords a biosensor by which to infer changes in the embedded enzyme’s activity.

**Fig. 4.**
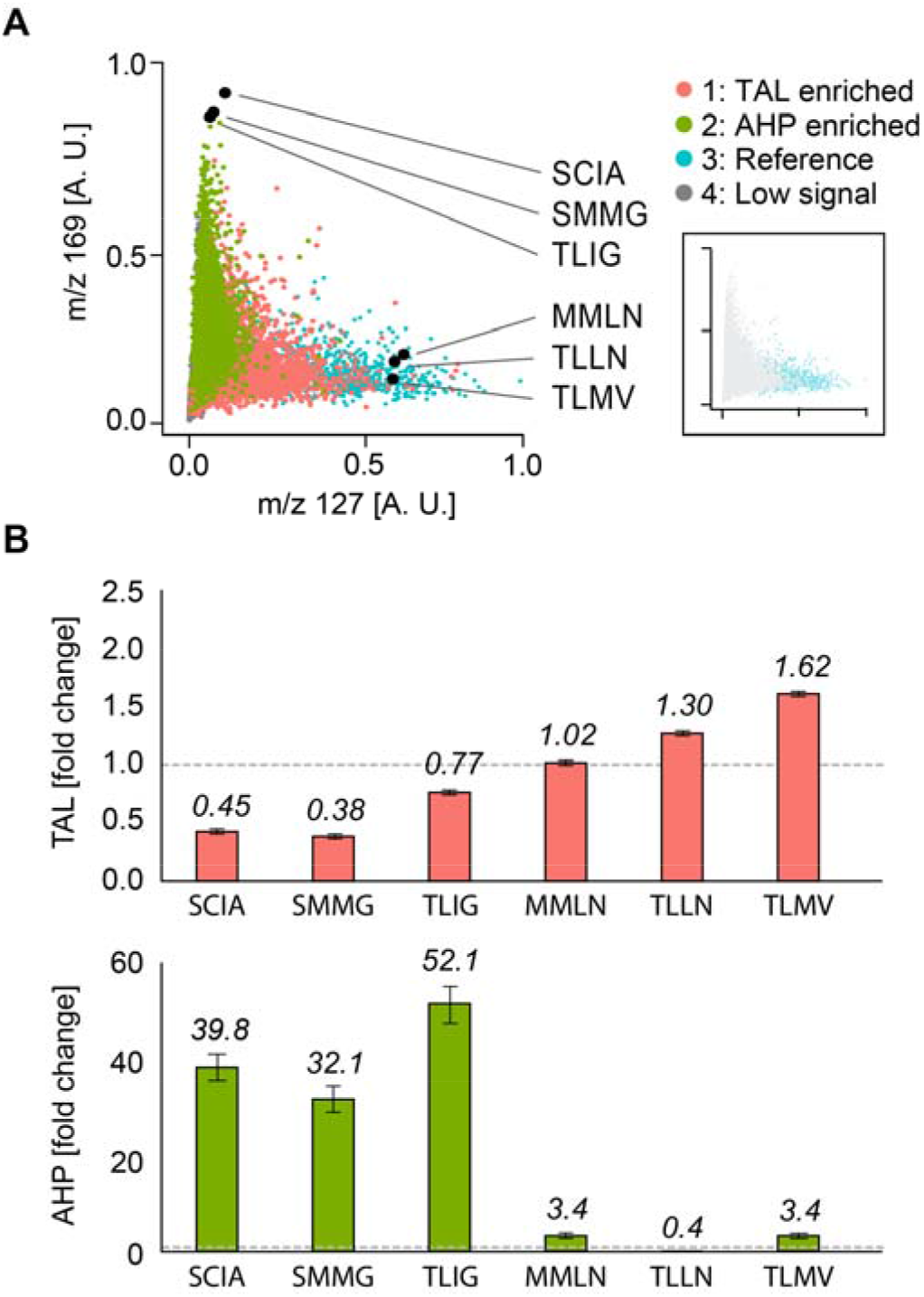
μMS screen of g2ps1 mutant library for increased production of TAL or AHP. **a**, Normalized m/z intensity for m/z 127 (TAL) versus 169 (AHP). Data points are colored based on clusters identified from UMAP analysis in Figure 2. Inset shows reference strain samples. The highest producers for TAL and AHP were isolated, sequenced, and confirmed in bulk analysis (black circles). **b**, TAL and AHP for selected bulk cultures as fold change over wild type (TLML).

## Discussion

Metabolomic biosensing allows inference of product formation even without direct product detection because the enzyme’s activity perturbs the host cell’s metabolite profile. Provided such perturbations are distinct for each activity, all activities generated by a mutant library can be detected, similar to the “compressed sensing” of smell sensation^19^. Such indirect sensing has beneficial features for enzyme engineering. In addition to providing a universal readout of product formation, the metabolites used to generate the UMAP can be selected to achieve the best cluster differentiation, even if the direct enzyme products are not included because they are difficult to detect due to their chemical or ionization properties. Furthermore, it should allow detection of all possible products, since each activity should perturb the host metabolite profile in unique ways and, thus, generate distinct clusters in the UMAP. By subsampling the clusters, unexpected products can be discovered. While we have utilized this analysis approach with yeast cells and MALDI-MS, it should apply to other bioproduction systems, including cell-free extracts, bacteria, and mammalian cells. Other readout modalities that provide sensitive, multiplexed detection of metabolites should be applicable, including other forms of mass spectrometry^20^ and multiparametric spectroscopy^21^. Lastly, while we have focused on an enzyme library for its simplicity, the approach should apply to any engineering that perturbs host cell metabolism, including genetic circuits and biosynthetic pathways.

## Acknowledgements

This work was supported by the ChanZuckerberg Biohub and the National Institutes of Health(NIH) (Grant Nos. R01-EB019453-01 DP2-AR068129-01 and R01-HG008978-01). E. M. P. was supported by NIH T32 EB005582. R. T. K. thanks to the NSF grant CHE-1904146

## Author Contributions

L.X. and A.R.A. conceptualized the project. L.X. designed and performed the experiments. L.X., KC.C., L.L., C.M.P., & H.S.P were involved in selecting, designing, and constructing the enzyme library. KC.C. prepared the library. L.X. prepared and processed the custom substrate and MALDI MS. N.T. & D.S.C provided access the Bruker MALDI and assisted in data acquisition. E.M.P. & R.T.K. assisted in the validation of selected mutants. L.X., KC.C, E.M.P., C.M., & A.R.A. analyzed and interpreted the obtained data. L.X., KC.C, C.M., & A.R.A. wrote the manuscript. All authors read and agreed to the final work.

## Methods

Methods, including statements of data availability and any associated accession codes and references, are available at https://doi.org/XXXXXXXXXX.

