## Supplemental for "Mapping enzyme catalysis with metabolomic biosensing"

**Supplementary figures and tables**

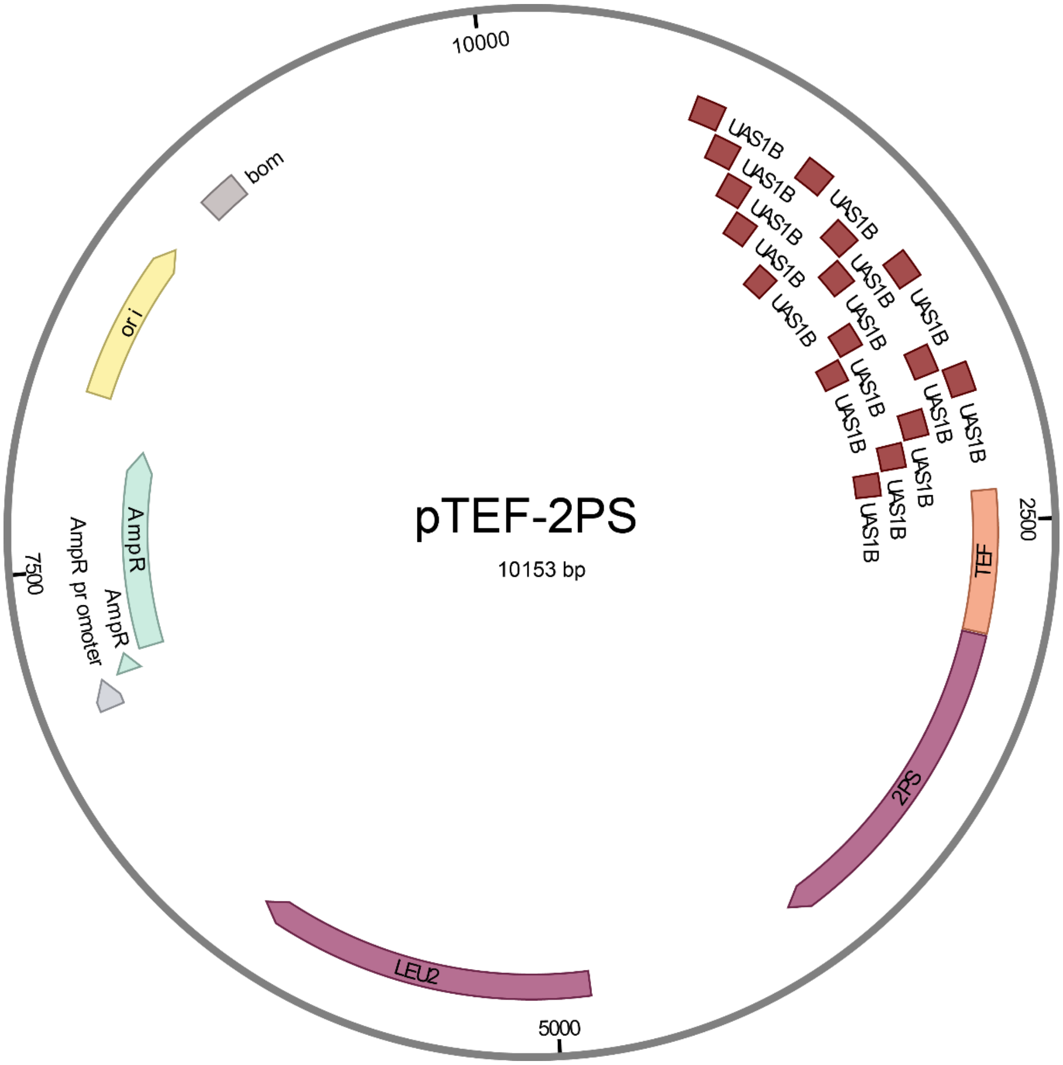


**Fig. S1 | Construct map of the plasmid pTEF-2PS.**

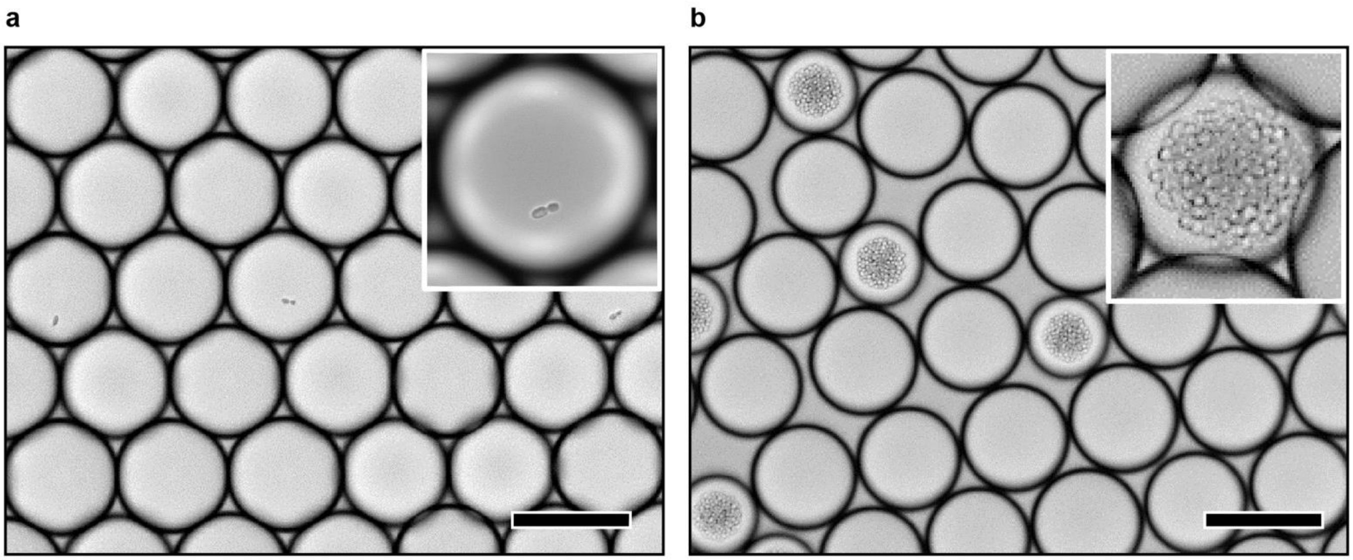


**Fig. S2 | Culture of *Y. lipolytica* Po1g strain in droplets. a,** Single cells are encapsulated in water-in-fluorocarbon oil droplets containing culture medium following a Poisson distribution. **b,** Single cells grown into isogenic microcolonies after a week of incubation. Scale bar: 100 µm.

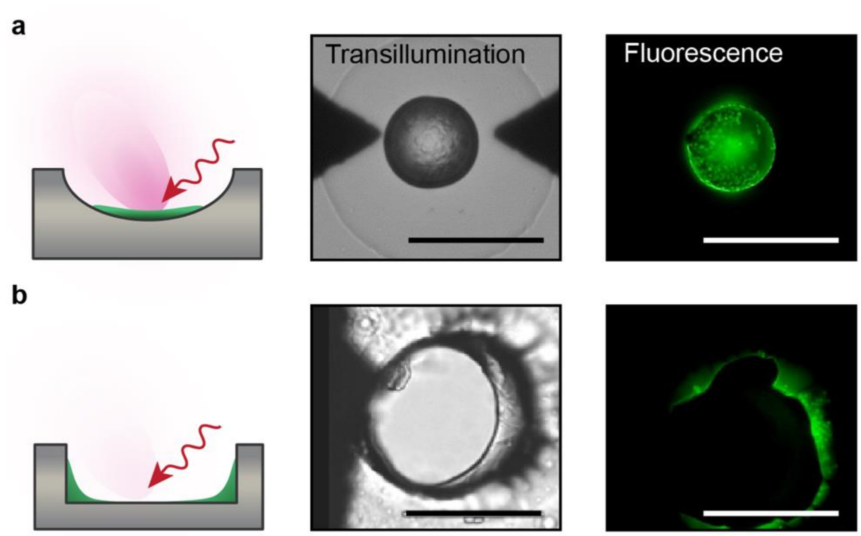


**Fig. S3 | Well shape affects colony desiccation and data quality in µMS. a,** Rounded bottom wells concentrate desiccated colonies to the center, facilitating laser ionization and yielding better, more reproducible measurements. **b,** Cylindrical wells concentrate material on the edges, where laser ionization is occluded. Scale bar for all images: 50 µm.

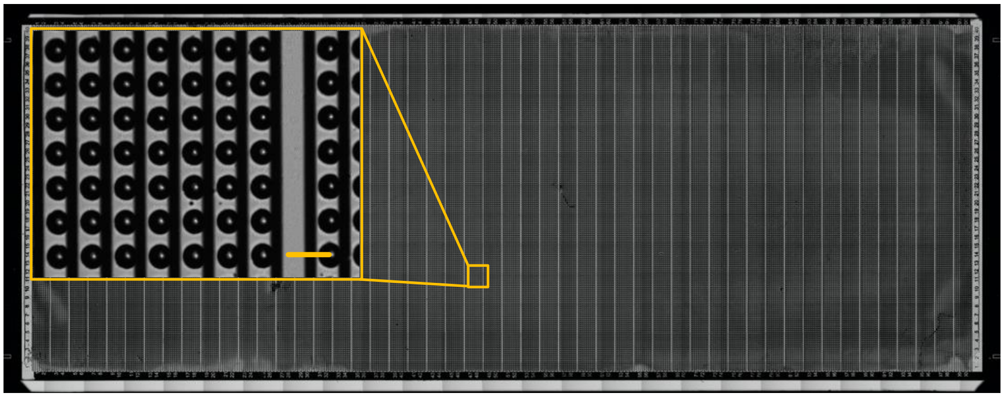

**Fig. S4 | 100k μMS well arrays.** The high-density substrate contains 100,000 round-bottomed wells and trapping electrodes on a 25-by-75 mm slide. Magnified yellow box shows individual wells. Scale bar: 150 μm.

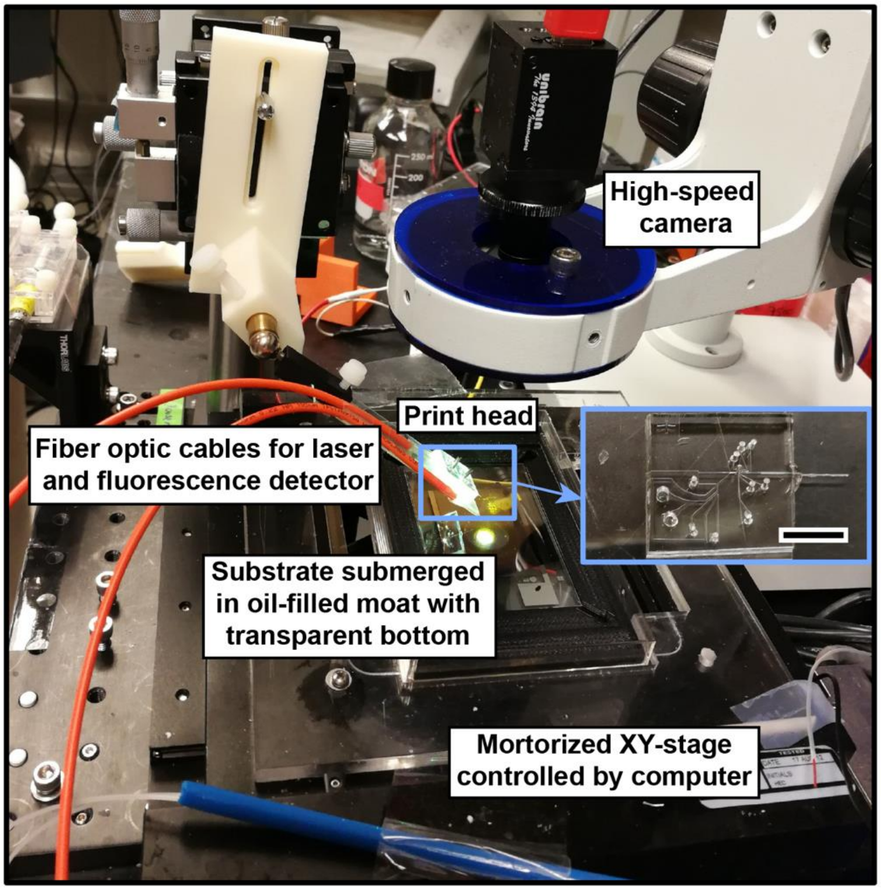


**Fig. S5 | Printed droplet microfluidic instrument.** Photograph of the PDM printing station built on an inverted microscope and comprising a mechanical stage for automated substrate positioning. The print head is attached to a fixed arm above the substrate and contains the fiber optics, electronics, and fluidics (blue inset) for droplet dispensing. Scale bar: 1 cm.

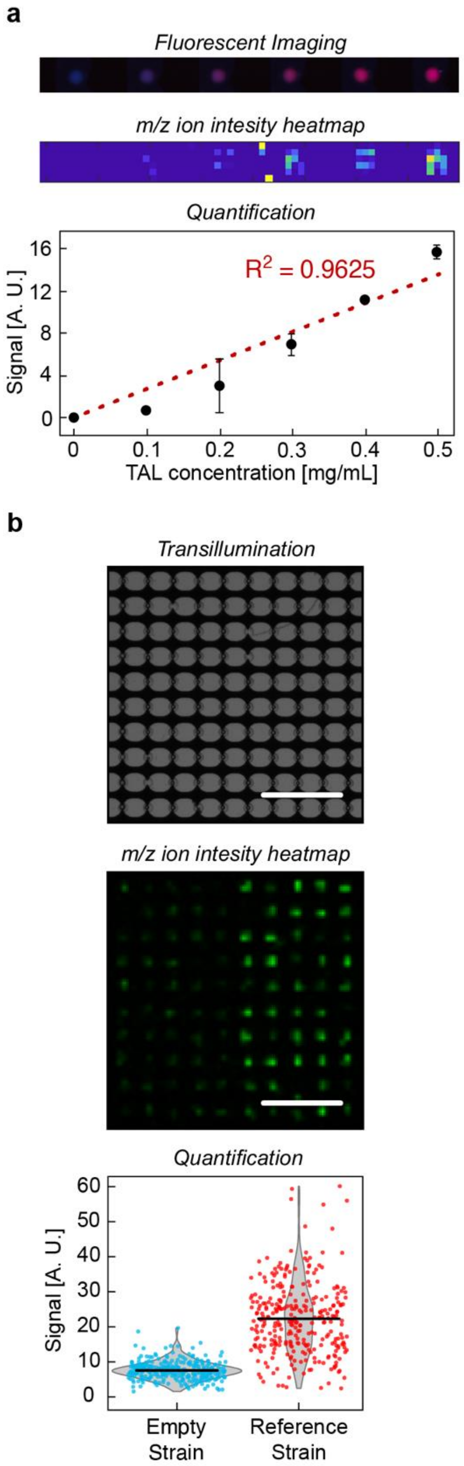


**Fig. S6 | Quantification test of µMS for printed droplet arrays. a,** Titration series of triacetic acid lactone (TAL). *Top*: Fluorescence imaging of printed droplets containing TAL and proportional amounts of Dextran-labeled dye. *Middle*: Signal intensity heatmap for TAL's 127 mass-to-charge (m/z) signal. *Bottom*: Concentration versus signal plot showing a linear response. Error bars represent coefficient of variation from the quantification. **b**, Test assay comparing empty vector and wild type enzyme controls. *Top*: High-magnification picture of well array with printed *Y. lipolytica* colonies. Scale bar: 1 mm. *Middle*: Resultant TAL m/z signal intensity heatmap of a printed population of either empty-vector or G2PS1-expressing *Y. lipolytica* cells.  Scale bar: 1 mm. *Bottom*: Scatter plot quantifying TAL level for 300 empty-vector and 300 G2PS1-expressing cells. Black bar represents population mean.

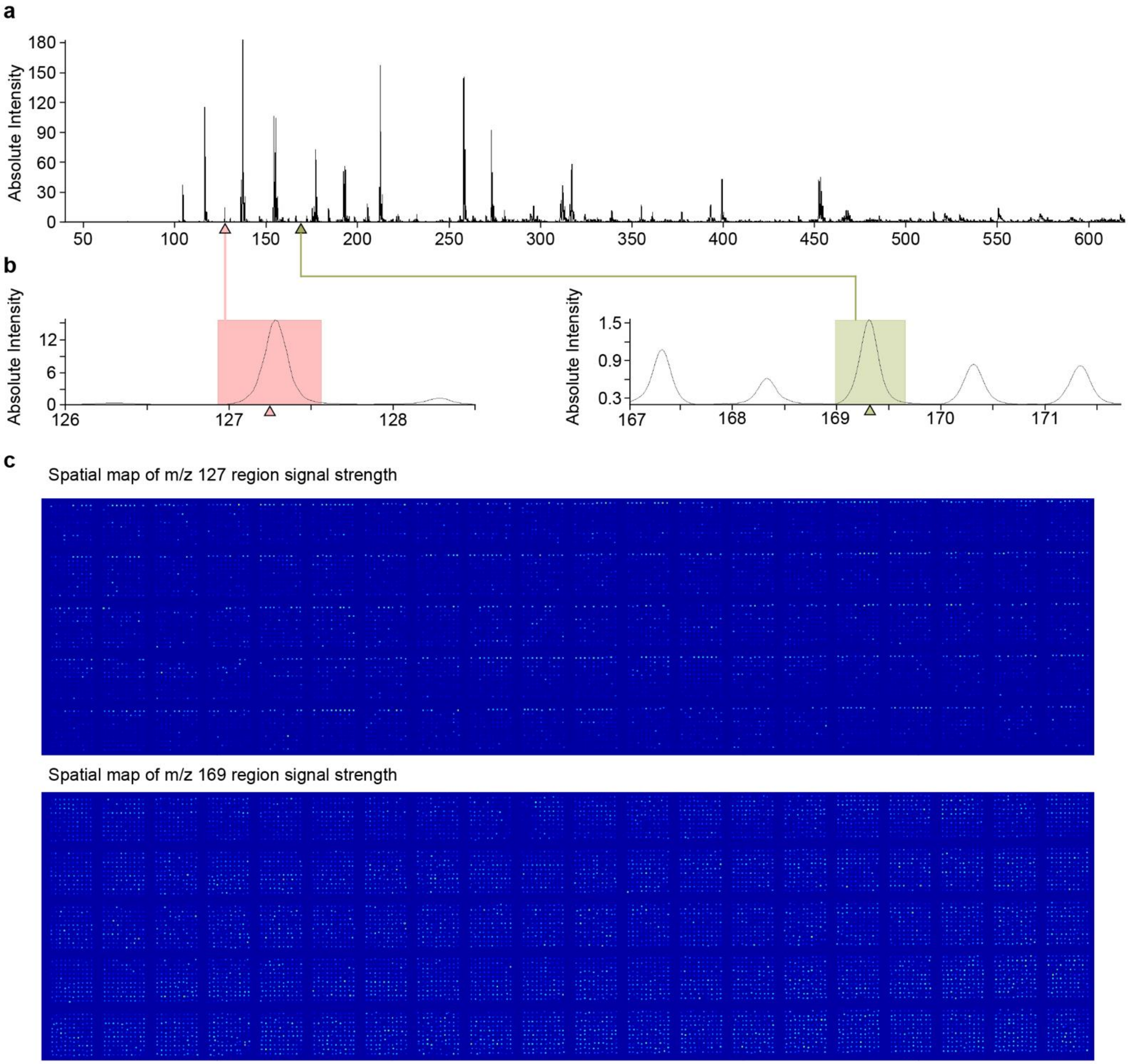


**Fig. S7 | Spectral imaging of μMS slide. a**, Sum of all mass spectra signals from all 10,000 wells after total ion normalization visualized with Bruker SCiLS Lab software. **b,** By selecting any m/z region within the range of 30 to 630, targeted metabolites can be interrogated for all 10,000 wells. The red box indicates the peak for TAL (127) and the green for AHP (169). **c**, Ion signal intensity can also be visualized across wells as an image for TAL (upper) and AHP (lower). The top row for each block of 100 contains a high TAL production reference strain.

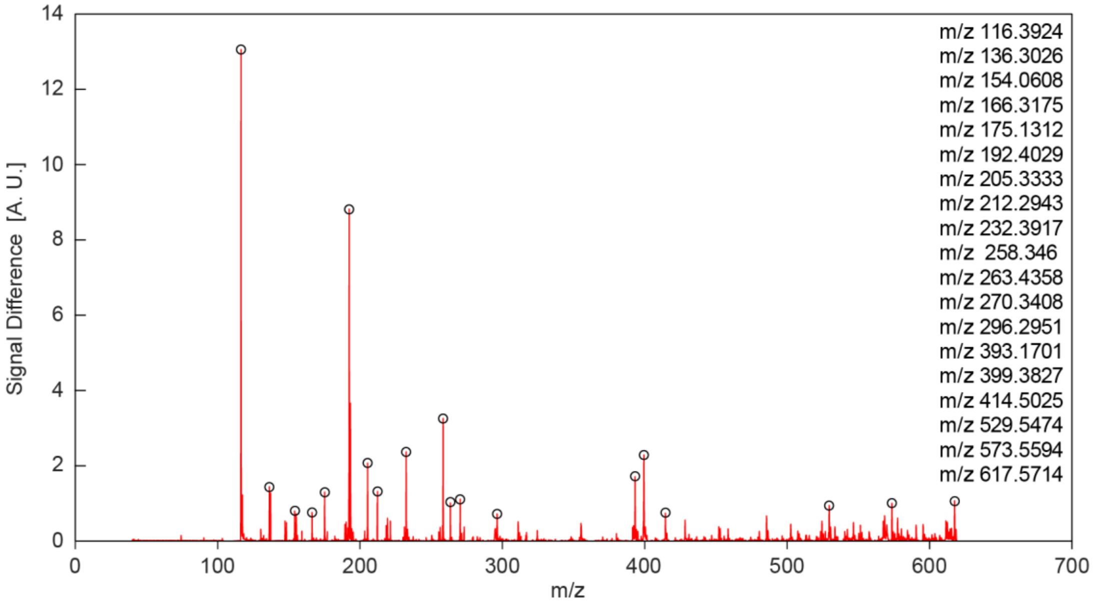


**Fig. S8 | m/z peaks used for 2D UMAP analysis of main text.** The top 19 brightest spectrum peaks for the signal sum from all wells of library subtracting all wells of reference are used to generate the UMAP.

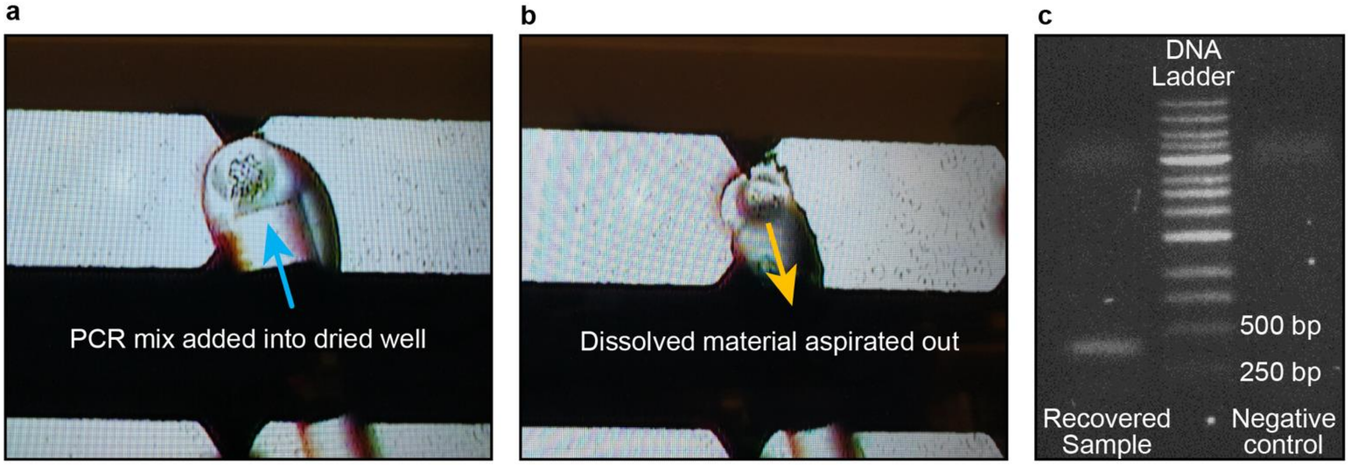


**Fig. S9 | Recovery of variant gene sequence after MALDI MS ion imaging. a,** a glass micropipette is used to add a water droplet to the dried yeast in the wells and then **b,** the water droplet containing yeast cells is withdrawn and transferred to a PCR tube for **c,** PCR of the mutation site (380 bp) and sanger sequencing.

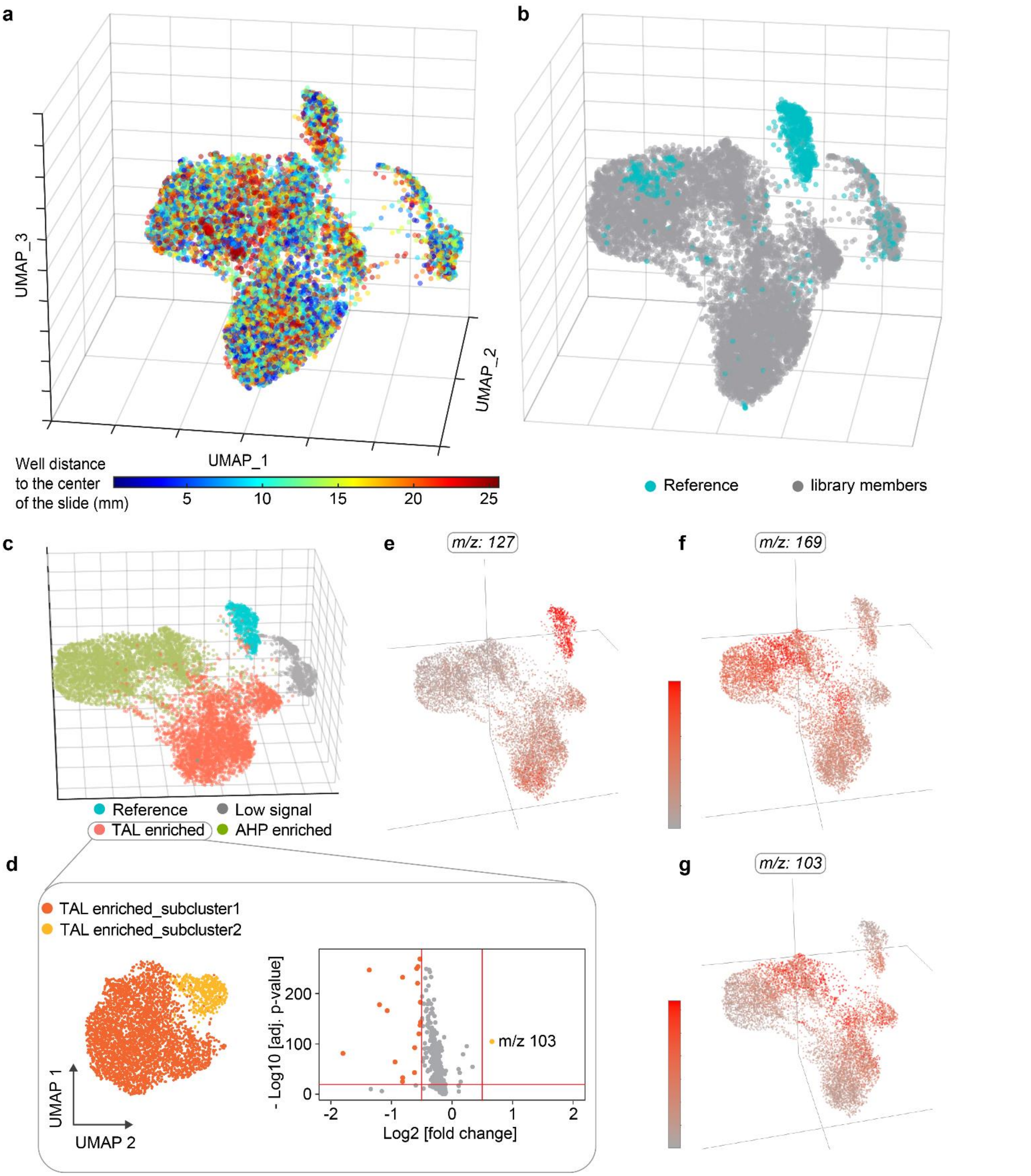


**Fig. S10 | Three dimensional UMAP based on peaks from the complete dataset. a,** UMAP clustering colored by well distance to the center of the slide. **b,** UMAP clustering with reference strain wells colored blue; the 3D UMAP including all m/z peaks does not group all reference points to a single cluster, whereas for the top 19 peaks they're localized (Figure 3C). **c,** 3D UMAP and **d,** 2D UMAP and volcano plot for just the TAL cluster. The 2D TAL UMAP demonstrates a novel cluster in which the peak m/z = 103 is highly differentially expressed, corresponding to a metabolite of unknown identity. 3D UMAP heatmaps for specific metabolites for **e,** m/z 127 (TAL), **f,** m/z 169 (AHP) and **g,** m/z 103 (unknown metabolite). Metabolite 103 appears to be highly expressed on the upper right ends of the TAL and AHP lobes, and may be related to an unknown activity of the enzyme.

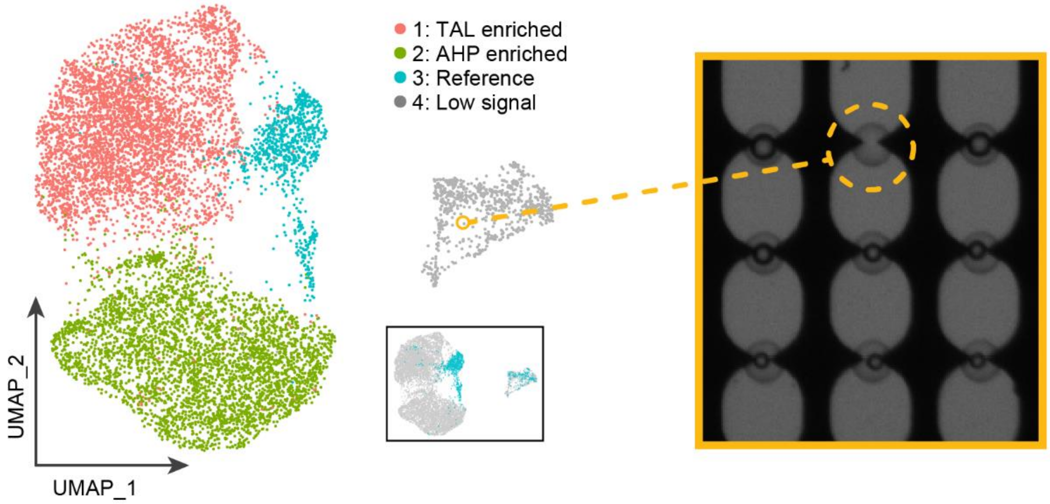


**Fig. S11 | Low signal cluster of the UMAP corresponds to empty or near-empty wells. a,** Transillumination imaging of well samples clustered within the UMAP cluster full of low signal samples indicates misprints of droplets that lack cell material.

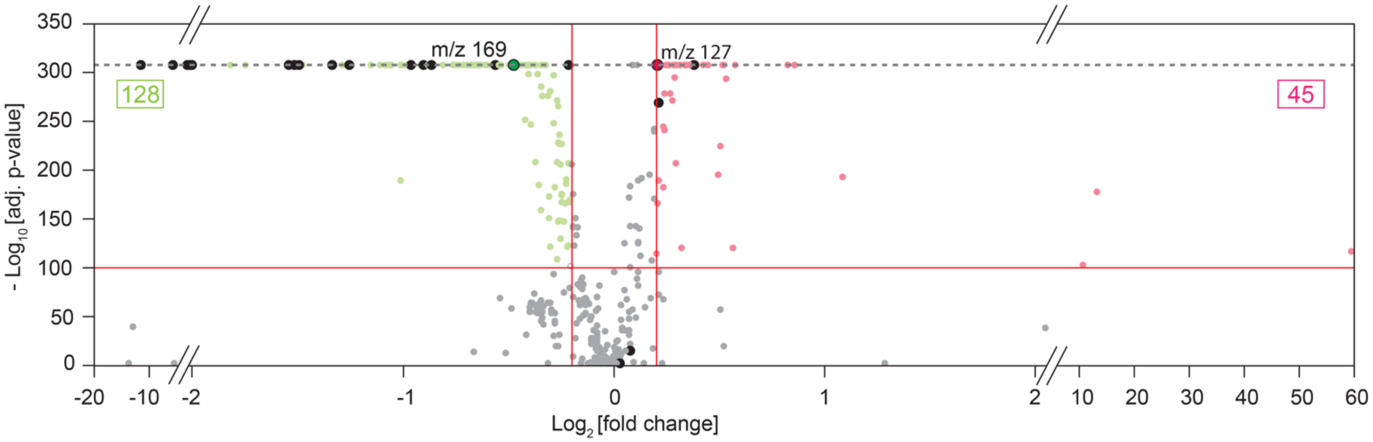


**Fig. S12 | Volcano plot between the TAL enriched and AHP enriched clusters.** The TAL cluster (red dots) is high in TAL (127) while the AHP (green dots) is high in AHP (169). Black dots indicate the nineteen m/z value that used to generate the UMAP. Other metabolites of unknown composition are also highly differentially expressed between these clusters and likely indicate the impact of the associated enzymes activity on the host cell metabolome. The horizontal red line represents the -Log10(adj. p-value) threshold at 100 and the vertical red lines indicate ±0.2 threshold values for Log2(fold change). Gray dash line indicates the smallest computational value in R for -Log10(adj. p-value) at an (adj. p-value) = 2.225074E-308.

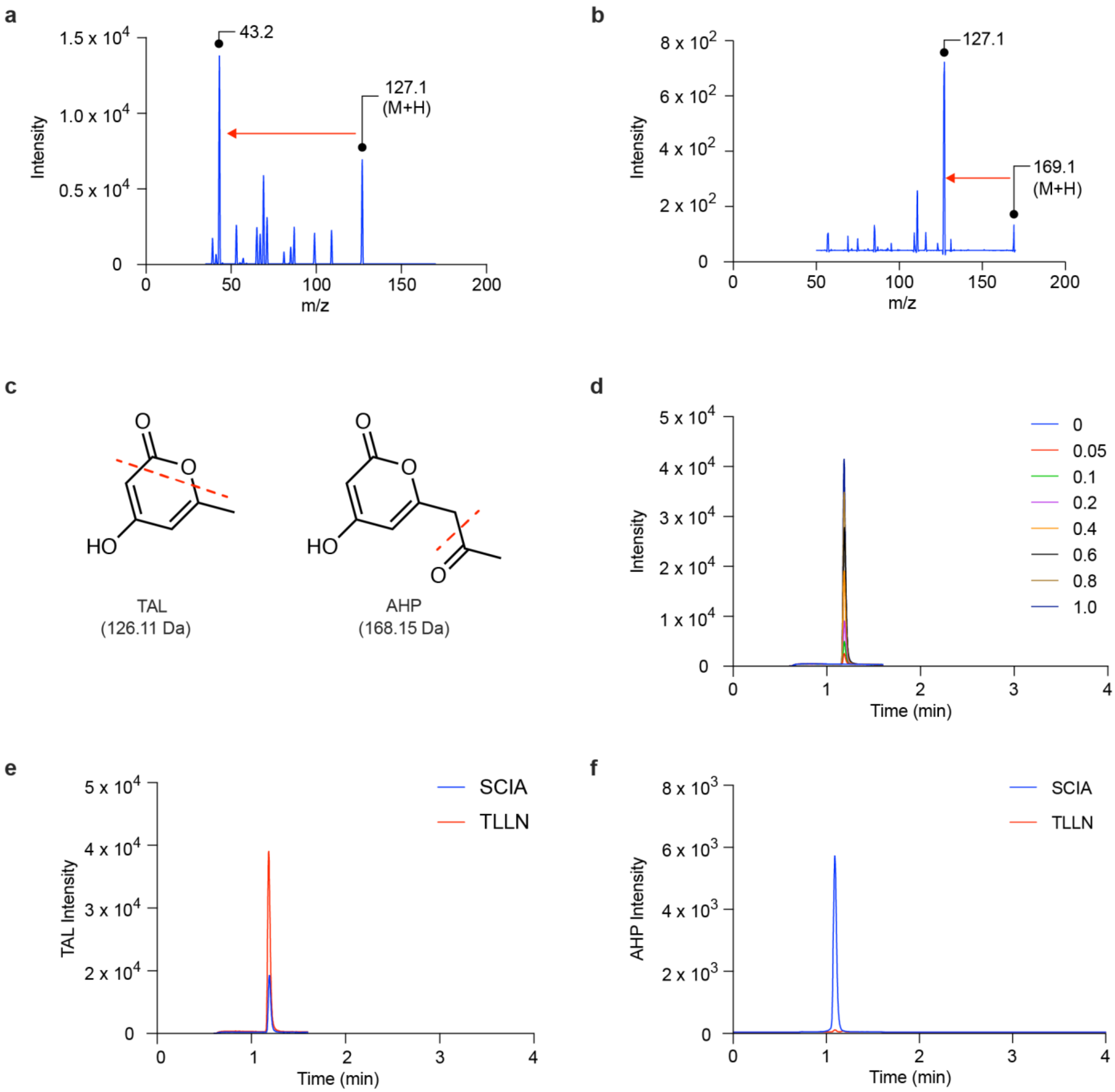


**Fig. S13 | HPLC-MS/MS of *Y. lipolytica* strains recovered from the wells. a,** Product ion mass spectrum of TAL showing intact ion (m/z: 127.1) and most abundant product ion (m/z: 43.2). **b,** Product ion mass spectrum of AHP showing intact ion (m/z: 169.1) and most abundant product ion (m/z: 127.1). Red arrows in **a** and **b** denote monitored mass transitions for HPLC-MS/MS analysis. **c,** Structures and masses of TAL and AHP, with proposed fragment pattern observed by MS/MS shown with dashed line. **d,** Overlaid chromatograms of HPLC-MS/MS for TAL calibration samples. Legend titles represent concentration of TAL in mg/mL. **e,** Overlaid TAL mass chromatograms for SCIA and TLLN mutants. **f,** Overlaid AHP mass chromatograms for SCIA and TLLN mutants. AHP abundance for TLLN production is too low to detect.

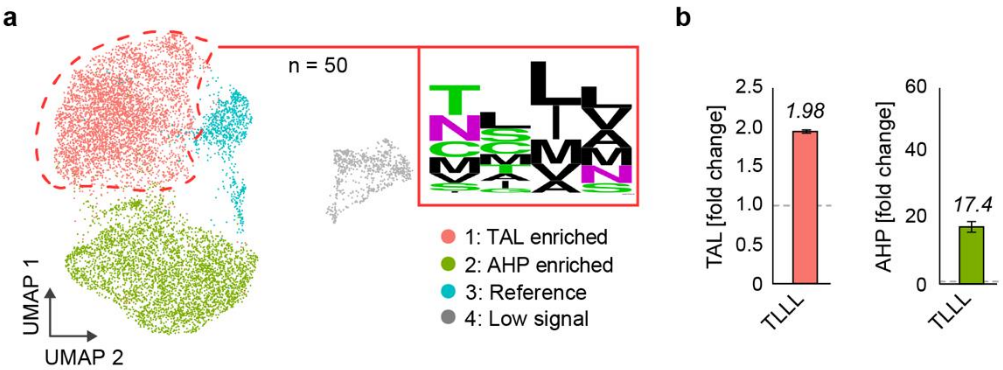


**Fig. S14 | Synthesis of hypothesized superior mutant for TAL production based on sequence consensus of TAL UMAP cluster. a,** ~50 mutants from the TAL enriched cluster are recovered and sequenced to generate a web logo. **b,** The predominant residue set, TLLL, has enhanced TAL and AHP production compared to wild type.

**Table S1 | Raw data of m/z peaks for each corresponding wells.** After total ion normalization, binning and averaging the signals of all wells, 399 m/z peaks that are three times above the background are selected (the 19 m/z peaks used for UMAP clustering are included). Wells that do not have signal intensity above the background threshold (43 wells) or that are abnormally high (17 wells) are removed, resulting a total of 9940 wells that proceed in the analysis.

**Table S2 | Fold change of m/z peaks and adjusted significant P value for the TAL cluster and AHP cluster using Wilcoxon Rank Sum test.**

**Table S3 | Sequences at the 4 active site positions (199 202 259 261) of the top screened mutations of g2ps1 enzyme for TAL and AHP respectively and their bulk production levels verified by HPLC tandem MS.**

**Table S4 | Sequences at the 4 active site positions (199 202 259 261) of mutations recovered from TAL and AHP clusters based on the UMAP analysis.**

**Table S5 | Plasmid sequences of ptef-2ps.**

**Video S1 | Three dimensional UMAP featured with types of colonies in wells as shown in Figure S10b.**

**Video S2 | Three dimensional UMAP as shown in Figure S10c.**
